# Investigating the temporal pattern of neuroimaging-based brain age estimation as a biomarker for Alzheimer’s Disease related neurodegeneration

**DOI:** 10.1101/2022.03.18.484935

**Authors:** Alexei Taylor, Fengqing Zhang, Xin Niu, Ashley Heywood, Jane Stocks, Gangyi Feng, Karteek Popuri, Mirza Faisal Beg, Lei Wang, the Alzheimer’s Disease Neuroimaging Initiative

## Abstract

Neuroimaging-based brain-age estimation via machine learning has emerged as an important new approach for studying brain aging. The difference between one’s estimated brain age and chronological age, the brain age gap (BAG), has been proposed as an Alzheimer’s Disease (AD) biomarker. However, most past studies on the BAG have been cross-sectional. Identifying how an individual’s BAG temporal pattern changes over time would enable improved prediction of clinical outcome based on neurophysiological changes and better understanding of AD progression. To fill this gap, our study conducted predictive modeling using large neuroimaging data with up to 8 years of follow-up to examine the temporal patterns of the BAG’s trajectory and how it varies by subject-level characteristics and disease status. To the best of our knowledge, this is the first effort to take a longitudinal approach to investigate the pattern and rate of change in BAG over time in individuals who progress from mild cognitive impairment (MCI) to clinical AD. Combining multimodal imaging data in a support vector regression model to estimate brain age yielded improved performance than single modality. Multilevel modeling results showed the BAG followed a linear increasing trajectory with a significantly faster rate in individuals with MCI who progressed to AD compared to cognitively normal or MCI individuals who did not progress. The dynamic changes in the BAG during AD progression were further moderated by gender and APOε4 carriership. Findings demonstrate the BAG as a potential biomarker for understanding individual specific temporal patterns related to AD progression.

## INTRODUCTION

Alzheimer’s Disease (AD) is the 6^th^ leading cause of death in the U.S., affecting 1 in 9, or 6.2 million Americans over the age of 65 as of 2021 [1]). Crucially, AD-related brain changes precede clinical symptoms, hampering efficacy of treatment [2–4]. Thus, exploring the trajectory of AD-related brain changes would likely lead to improved early detection through prediction of future outcomes. While aging in cognitively normal (CN) individuals ultimately leads to some structural brain atrophy [5], neurodegenerative diseases such as AD show rapid deviation from the normal aging trajectory, particularly with regard to gray matter [6]. To understand the temporal pattern of these AD-related deviations from normal aging, predictive models using large longitudinal neuroimaging datasets can be leveraged.

Recently, machine-learning based methods have been used to estimate a person’s brain age using neuroimaging data, allowing researchers to investigate the difference of this estimated brain age to the participants chronological age [7–11]. Referred to as the brain age gap (BAG), it has been used to examine brain aging in major depressive disorder [12], Parkinson’s disease [13], schizophrenia [14], as well as non-disease related differences in cognitive maintenance [15] and lifestyle behaviors [16]. In AD, a positive BAG value (where the estimated brain age is greater than the chronological age) has been associated with increased risk of dementia onset using unimodal [7,10,17] and multimodal neuroimaging measures [18], indicating its potential as a personalized AD biomarker. Furthermore, individuals at an intermediate stage of AD-related brain changes, known as mild cognitive impairment (MCI), exhibit spatially distinct patterns of gray matter loss consistent with AD related neuropathology [19]. Studies showing significantly greater BAG at later stages of AD and association with clinical symptom severity reflect this [20,21], suggesting the BAG may follow a nonlinear trajectory. While pinpointing the exact biological differences quantified by the BAG remains a challenge, its association with brain diseases warrants further investigation.

However, literature relating the BAG to AD progression is sparse. Most studies used cross-sectional designs [18], which are inadequate in describing trajectories and do not simultaneously consider intra- (i.e., within subject neurophysiological changes) or inter- (i.e., between subject variance) individual differences [22,23]. A limited number of longitudinal studies compared study timepoints only to baseline assessment [20,21], but this approach does not fully investigate potential non-linear trajectories of the BAG. For example, these studies found BAG scores compared between baseline and a follow up time in several years were significantly greater in progressive MCI and AD individuals compared to earlier or stable groups, though whether this remains significant after considering variation across multiple timepoints is unclear. Identifying how an individual’s BAG temporal pattern changes over time (i.e. linear or non-linear) would enable improved prediction of clinical outcome based on neurophysiological changes.

Together, this data requires consideration of the covariance structure among repeated measures across individuals and groups must be considered to account for the heterogeneity common in longitudinal datasets and in particular, AD patient data [24,25]. The appropriate analytical design would then suggest either a linear or non-linear fit to the data. Extant literature on brain and cognitive aging have shown both linear and non-linear changes across age groups, brain structures, and cognitive abilities [26,27]. One other study of note examined longitudinal patterns of brain atrophy between CN and MCI subjects using magnetic resonance imaging (MRI) data and found accelerating brain aging for older CN and MCI individuals, though the results were based on a small sample size and did not distinguish between those with stable or progressive diagnoses, and in addition did not explore the moderating effects of covariates. [28].

Choice of neuroimaging modality has also been homogenous in the BAG literature. Typically, structural T1-weighted images captured using MRI have been used to estimate BAG. While MRI derived cortical thickness inform structural brain changes and have been used in AD prediction [29], fluorodeoxyglucose positron emission tomography (FDG-PET) provides complementary functional information through an indirect measure of metabolic function via glucose consumption [30,31]. Studies using this modality have shown neuronal dysfunction can precede gray matter atrophy [32,33], and has been used to measure neurodegeneration [34]. However, FDG-PET’s use in BAG prediction, and particularly understanding its temporal pattern, has not been extensively studied [35], despite its utility in predicting AD [36]. Importantly, it has been suggested to show more consistent functional changes at earlier stages of AD compared to MRI [31], making it an ideal modality for tracking biological changes preceding clinical symptoms of AD. This presents an opportunity to utilize both modalities for understanding the BAG trajectory. In literature unrelated to brain age prediction, the combination of MRI and FDG-PET improves discrimination prediction performance of AD [37] and MCI [38]. As the use of FDG-PET has not been fully explored in brain age prediction, and given that multiple modalities improves performance of brain age prediction models [9,18], the use of both FDG-PET and MRI images is warranted.

In the current study, we tested the hypothesis that the BAG’s temporal pattern would be non-linear, increasing at a faster rate in individuals who were initially diagnosed as MCI but progressed to AD, vs. those who remained diagnosed as CN or MCI. To study the temporal pattern, we utilized longitudinal data from the Alzheimer’s Disease Neuroimaging Initiative (ADNI) (http://adni.loni.usc.edu/) from individuals with up to 8 years of follow-up. Further, as BAG has been shown to be moderated by factors such as sex and/or gender [39] and carriership of the apolipoprotein E ε4 (APOEε4) allele [21], we also examined their influence on the BAG temporal patterns. To the best of our knowledge, this is the first longitudinal study to investigate the pattern (linear vs. non-linear) and rate of change in BAG over time in individuals who progress from MCI to clinical AD. With further validation, BAG as a biomarker for AD has clinical application in assessing brain health and providing insight into a patient’s brain age trajectory [11].

## METHODS

### Participants

Data were downloaded from the Alzheimer’s Disease Neuroimaging Initiative (ADNI) database that included ADNI1, ADNI-GO, ADNI2, and ADNI3 (adni.loni.usc.edu/). The primary goal of ADNI has been to test whether neuroimaging, clinical, neuropsychological, or other biological markers can be combined to measure the progression of MCI and early AD.

ADNI participants were stratified into three groups: stable CN (sCN), stable MCI (sMCI), or progressive MCI (pMCI). Stable individuals maintained the same diagnosis, either CN or MCI, for the duration of ADNI. Progressive individuals changed from a baseline assessment of MCI diagnosis to an assessment diagnosis of AD for the duration of ADNI without reverting to MCI. The total number of assessment time points varied across subjects. Participants were only included if they had two or more assessments (including baseline) and had both MRI and PET scans. The final study sample included 168 sCN, 217 sMCI, and 108 pMCI individuals. Reported gender (male or female), age, years of education, race, mini-mental state exam (MMSE) score, and APOEε4 allele carriership were provided from the ADNI dataset.

Demographic and clinical measurements were documented and available for each assessment point for all individuals. APOEε4 carriership was determined at the initial enrollment of that individual. Baseline data from participants in the three groups were compared in reported gender, age, years of education, race, MMSE score, and APOEε4 carriership. One-way ANOVA between groups was carried out for variables age and education, while MMSE scores were compared using a permutation-based ANOVA. Categorical data of gender and APOEε4 were compared using Chi-Square tests. The variable race was analyzed using Fisher’s Exact test.

### Imaging Data

Structural MRI and FDG-PET imaging measures used for brain age estimation were provided by Popuri et al (2018) [40][40]. In that study, structural MRI data were processed with FreeSurfer v5.3 [41], generating 85 cortical and subcortical gray matter regions of interest (ROI) (Table S1). In the present study, structural MRI mean cortical thickness values were used in subsequent analysis for brain age estimation. Following co-registration of FDG-PET with structural MRI, Popuri et al. (2018) parcellated each ROI into equal-size patches to mask the FDG-PET data for improved localization. Patch-wise standardized uptake value ratios (SUVRs) of FDG-PET were provided for a total of 16 patch-size levels: 100 voxels per patch, 150, 200, 250, 300, 350, 400, 450, 500, 1000, 1500, 2000, 3000, 4000, 5000, and 10,000 (Popuri et al., 2018). To reduce computation time, we evaluated 5 patch-size levels (500, 1000, 2000, 5000, and 10,000) in the training step of the brain age estimation analysis (explained in the following section below) and selected the patch size of 2000 voxels/patch based on it returning the lowest root mean squared error (Table S2). This 2000 voxels/patch size resulted in 66 and 65 left and right hemisphere MRI features, respectively, and 343 FDG-PET features across cortical and sub-cortical structures. Subsequent brain age estimation analyses were based on SUVRs at this patch size.

### Brain Age Estimation

Data preparation and analysis were carried out using *R* (v3.6.2, R Core Team, 2019). Brain age estimation was accomplished using a support vector regression (SVR) model (*e1071* package (v1.7-3) within *R*) [42], given that the dependent variable of age was continuous, and that this method has been frequently used in brain age prediction literature [7,9,18,43,44]. Briefly, SVR transforms a training set of data into high-dimensional feature space and attempts to fit the data by not penalizing error less than a threshold while minimizing model complexity through its hyperparameters. In order to observe deviations from healthy brains, the SVR was trained on baseline sCN data, then applied to data at all other time points in all individuals.

SVR model selection procedures are visualized in Figure 1. For this procedure, we included an additional 59 sCN individuals who had only baseline neuroimaging data (and thus were not included in the longitudinal analysis) for a total sample of *N* = 227. This sample was randomly split (without replacement) into a training set (75% of the data) and test set (remaining 25%), where all subjects had neuroimaging data at 5 patch-size levels (500, 1000, 2000, 5000, 10000 voxels/patch). In short, a different SVR trained with baseline neuroimaging data at each patch size was used to predict age, and the resulting prediction performances were compared to select the best model and patch size.

**Figure 1.**
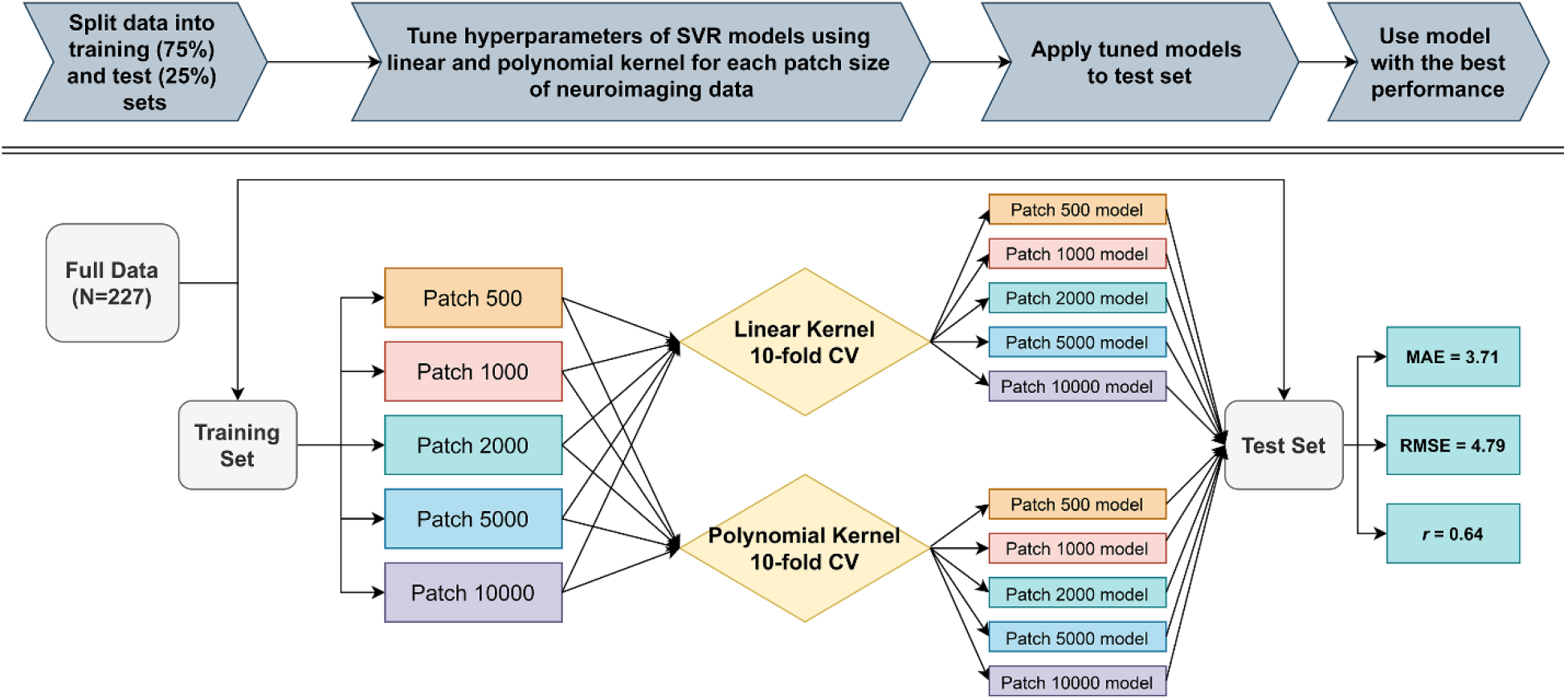
Model selection pipeline. 227 stable cognitively normal (sCN) participants were split into a training set (75% of the sample) and test set (25%), where each participant had baseline neuroimaging data at 5 different patch sizes (500, 1000, 2000, 5000, 10000 voxels/patch). A separate support vector regression (SVR) was trained and tested with each patch size to predict age. Hyperparameters of each SVR were optimized using 10-fold cross-validation with either linear or polynomial kernel. Each optimized SVR of a specific patch size then predicted age on the test set, where the best performing model was chosen as the one with the lowest root mean squared error (RMSE). This model along with neuroimaging data at the associated patch size were then used for all future analyses. Mean absolute error (MAE) and Pearson correlation (*r*) were also calculated for comparison to past studies.

Using either a linear or polynomial (to identify non-linear relationships) kernel, hyperparameters (i.e., cost parameter for both kernels, and gamma, degree, and constant term for polynomial kernel) of each SVR were optimized using 10-fold cross-validation. Briefly, this commonly used procedure splits the training data into 10 partitions, where 9 partitions are used to fit the model, and the remaining partition is used as a validation set to estimate the sample error. This is repeated 10 times such that each partition is used as a validation set once and the sample error is averaged across validation sets. Crucially, only data from the validation sample is used to estimate error during this procedure. Following 10-fold cross-validation, each SVR then predicted age on the test set, which contained participants unseen during model training to produce a valid estimate of prediction performance. Thus, the best performing model was chosen as the one with the lowest root mean squared error (RMSE). This model and the neuroimaging data from the associated patch size were then used for all future analyses. Additionally, performance metrics mean absolute error (MAE) and Pearson correlation coefficient (*r*) were computed. The estimation of brain age by the best performing model is subsequently referred to as estimated brain age.

This best performing model was then applied to all individuals in each group (i.e. sCN, sMCI, and pMCI) to obtain individualized estimated brain ages at each available assessment time point (not just baseline). The BAG was calculated as the difference between the individual’s estimated brain age and actual chronological age, where a positive BAG represents greater than expected brain aging. Similar procedures of SVR model training and testing were done using single modality MRI or FDG-PET data, but these did not yield superior prediction performance compared to the model using combined modalities.

Finally, a well-documented observation in brain age prediction is the bias towards overestimating the BAG for younger subjects and underestimating it for older subjects, likely due to regression to the mean [45–47]. To overcome this bias, BAGs should be further corrected for the confounding effects of chronological age. We evaluated a linear regression model on sCN (see [45] for further description of this procedure) and showed that this could successfully remove the bias (i.e., the expected age-regression in sCN is flat and centered around 0) (Figure 2B, left panel).

Notably, the bias is not eliminated in the sMCI and pMCI panels, suggesting especially for pMCI that the unbiased BAG is positive. It is important to note that correction for age-bias was not done during SVR training to avoid artificially boosting the model’s performance [48]. It was done in machine learning evaluation in subsequent analysis of BAG trajectory.

**Figure 2.**
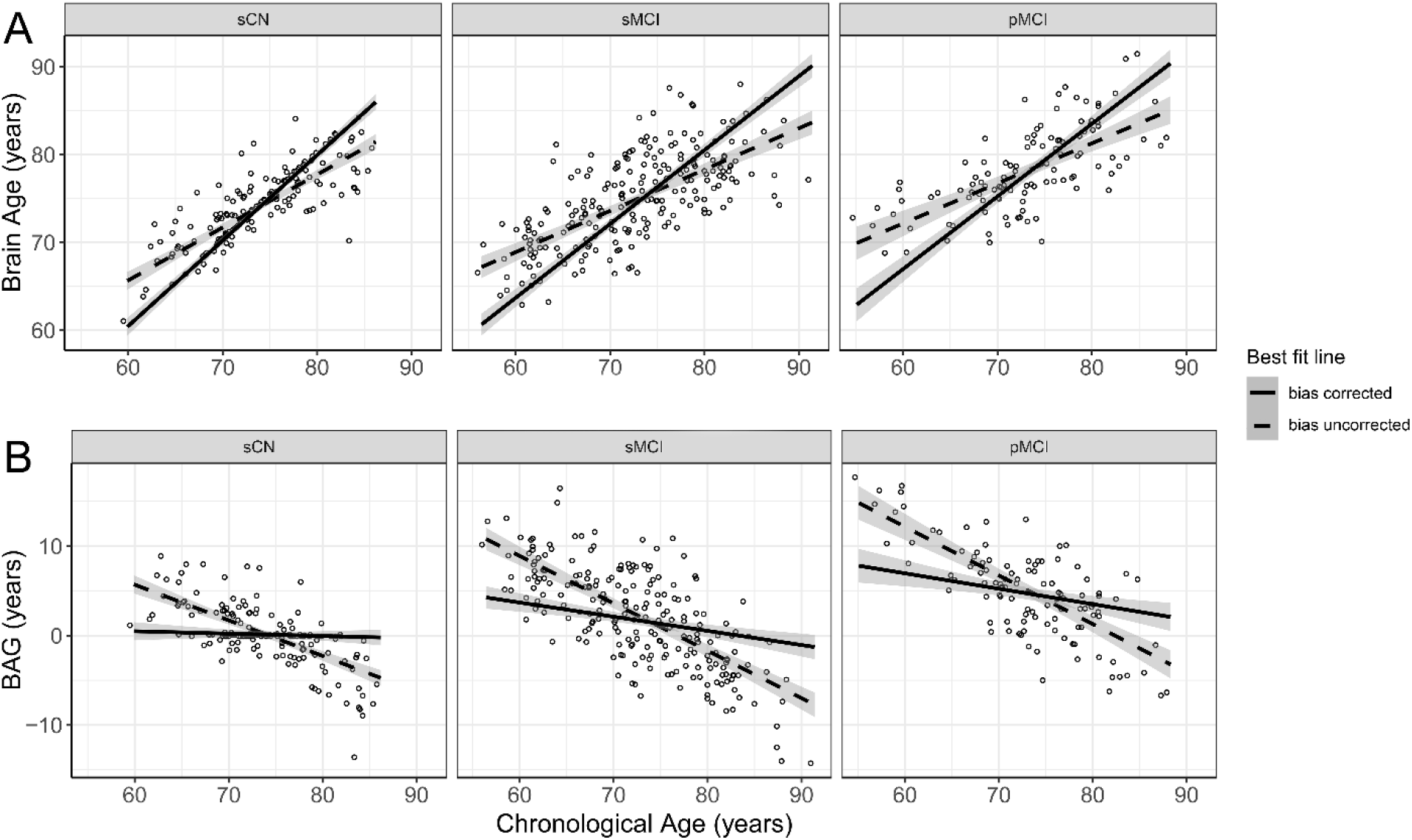
Comparison of best fit line for data before (dashed line) and after (complete line) bias correction is applied for each group. Groups are given as stable cognitively normal (sCN), stable mild cognitive impairment (sMCI), and progressive mild cognitive impairment (pMCI). For all plots, shaded area around each line indicates the 95% confidence interval. Individual points represent baseline data *prior* to bias correction. A: Baseline estimated brain age as a function of chronological age for all individuals in each group. B: Baseline brain age gap (BAG) as a function of chronological age for all individuals in each group.

### Feature visualization

To aid clinical interpretability, MRI and FDG-PET features were quantitatively labeled depending on their importance for SVR prediction performance. For instance, if the removal of one feature results in a model of greater MAE value compared to the MAE of a model using all features, that removed feature would be considered important for predicting sCN brain age. Thus, the importance of each feature was gauged by removing one feature at a time and calculating the resulting model’s MAE. Specifically, the ratio of MAE after removing a feature, over the original MAE was quantified, where a higher positive value indicated greater importance for sCN brain age prediction. Features were then min/max scaled, and any features that were divided into multiple patches (during preprocessing) were averaged together to create single overall importance values for each ROI. Top features were identified as being at least 1 standard deviation from the mean. Cortical surface features were visualized using SurfStat [49].

### Longitudinal analysis of BAG trajectory

Comparisons in BAG trajectory among groups were performed using multilevel modeling with the *lme4* package (v.1.1-21) in *R*. Multilevel modeling allows simultaneous quantification of inter- and intra-individual level patterns in the data, while taking advantage of the regression framework to examine effects of different covariates [50]. All models assessed the random effects of individuals by allowing their slopes and intercepts to vary across time. The time variable was defined as the assessment time point, coded as the number of months following baseline. Restricted maximum likelihood was used to estimate model parameters and to test the significance of random effects. The basic formation of multilevel models examined time as linear. Higher-order time effects were tested based on model selection criterion such as Akaike Information Criterion (AIC) and Bayesian Information Criterion (BIC). Following this determination, fixed effects of group type and its interaction term with time (either between sCN and pMCI, or sMCI and pMCI, where the pMCI factor was always compared against the sCN or sMCI reference factors) were added to the model. Additional covariates of gender and APOEε4 were tested separately as a fixed effect in the model to determine if they significantly moderated the effect of group type on the BAG trajectory.

### Analysis of pMCI BAG trajectory

To investigate whether the BAG trajectory pattern for pMCI individuals followed a non-linear pattern, piecewise linear segments centered around the assessment of AD diagnosis were used, where assessments from baseline until the assessment before the AD diagnosis was labeled preAD, and assessment time for and following AD diagnosis was labeled postAD. Only pMCI individuals with at least one assessment following AD diagnosis were included (*N*=47). Multilevel modeling was used estimate the rate of change in BAG for each segment, where segments preAD and postAD were used as separate fixed effects, and the random effects structure varied the slope and intercept of all pMCI subjects across preAD and postAD time segments.

To establish whether the difference in BAG trajectories (i.e., slopes) between segments was significant, an additional variable centered on the AD diagnosis was created to indicate the individual’s AD conversion status (assessments where pMCI individuals had converted to AD were coded as 1, otherwise they were coded as 0). This conversion variable was added as a fixed effect to the multilevel model to interact with the time effect. A significant positive estimate of this interaction would suggest the BAG slope following the diagnosis of AD was higher as compared to before the diagnosis, which would suggest that brain aging accelerated after diagnosis.

## RESULTS

### Participants

Age, gender, years of education, and race were not significantly different between the three groups (all *p* > 0.05), suggesting these variables would not confound later analysis. Differences in group MMSE (*p* < .001) and APOEε4 (*p* < .001) were significant between the three groups, though this was expected as CN individuals categorized as such due to lack of cognitive impairment and are less likely to be APOε4 carriers compared to AD individuals. Full demographic characteristics can be found in Table 1.

**Table 1.** Demographic information for each group. Stable cognitively normal (sCN) and stable mild cognitive impairment (sMCI) maintain the same diagnosis across assessments, while the group with progressive mild cognitive impairment (pMCI) progresses from MCI to at least a final diagnosis of Alzheimer’s Disease. Bold p-values are <.05.

|  | sCN (n=168) | sMCI (n=217) | pMCI (n=108) | *P-value* |
| --- | --- | --- | --- | --- |
| Female (%) | 72 ( 42.9%) | 86 (39.6%) | 42 (38.9%) | 0.752 |
| Mean Age (SD) | 74.2 (±5.69) | 73 (±7.55) | 73.5 (±7.14) | 0.259 |
| Mean MMSE (SD) | 29.1 (±1.13) | 27.9 (±1.75) | 27 (±1.66) | **<0.001** |
| APOEε4 carriers (%) | 42 (25%) | 83 (38.2%) | 71 (65.7%) | **<0.001** |
| Mean years of education (SD) | 16.5 (±2.82) | 15.9 (±2.83) | 16.2 (±2.61) | 0.166 |
| % White | 91.1% | 91.7% | 96.3% | 0.529 |

### SVR performance

The SVR was trained using baseline MRI and PET data from sCN individuals. The model with the best performance (i.e. lowest RMSE) was achieved using a linear kernel compared to polynomial. Estimation performance from the test set found an overall correlation of *r* = 0.64, MAE = 3.71 years, and RMSE = 4.79 years. These metrics are comparable to past studies using similar models based on CN data of similar age cohorts [8,10].

### Group BAG trajectory comparisons

Brain age prediction and bias correction models were then applied to all available assessment time points for each subject within each group. Mean BAG for the pMCI group was greater than the stable groups at every level of assessment (Figure 3), while a visual assessment shows sMCI and sCN maintained relatively stable BAG trajectories. Notably, the estimated variance in Figure 3 is minimal up until the final two assessment points, m84 and m96, where each group contained only several observations. Removal of these assessment points did not drastically change the significance or slopes of the following results.

**Figure 3.**
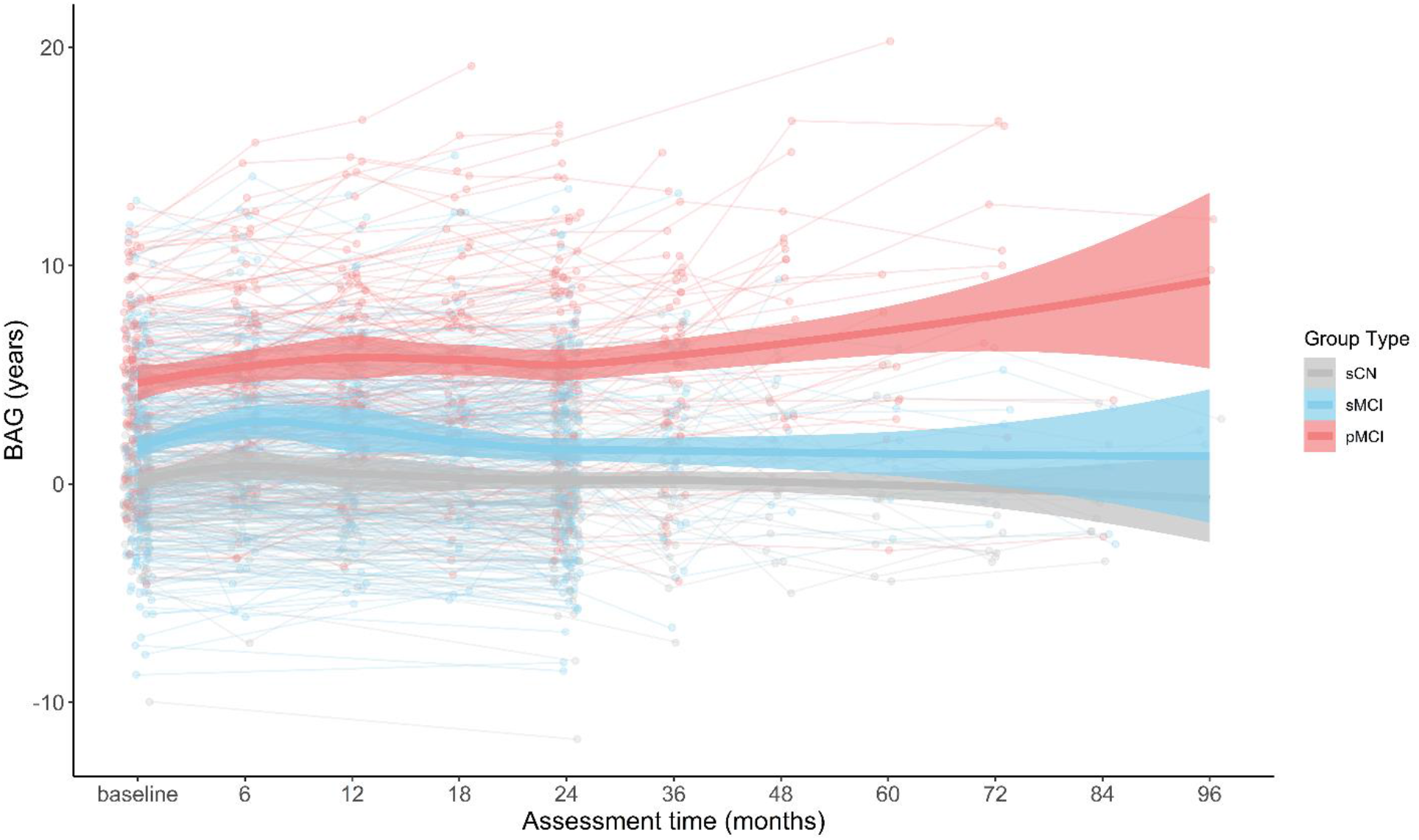
Individual and group brain age gap (BAG) values at each assessment month following baseline. Thick lines represent BAG of all subjects for each group across assessments, fitted by loess curve. Shaded area along each curve indicates the 95% confidence interval. Individual dots and lines represent individual subjects for each group. Groups are given as stable cognitively normal (sCN), stable mild cognitive impairment (sMCI), and progressive mild cognitive impairment (pMCI). Assessment time represents the months following the baseline (initial) assessment.

Multilevel modeling was used to assess the temporal trajectory of BAG over time (linear vs. nonlinear) across all individuals regardless of their group type. Given the large amount of variance among subjects in each group, a random effects structure varying subjects over assessment time was necessary to improve model fit (*X^2^*(2) = 212.8, *p* < .001). The pattern of change in BAG for all individuals was found to be linear over time (b =.090, *p* < .001). Inclusion of the fixed and random effects of quadratic time did not significantly improve the model fit.

In addition, we examined whether the rate of change in BAG over time was different for each group by including the 2-way interaction between assessment and group type. Comparing pMCI against the stable groups found the BAG of the pMCI group increased at a faster rate than the sCN group (*b* = .350, *p* < .001), and sMCI group (*b* = .310, *p* < .001). Results for all models are reported in Table 2 for sCN, and Table 3 for sMCI.

**Table 2.** Multilevel modeling results for group type stable cognitively normal (sCN) compared to progressive mild cognitive impairment (pMCI). Bold p-values are <.05

|  | Estimate | Std. Error | CI (95%) | Statistic | P-value |
| --- | --- | --- | --- | --- | --- |
| **Interaction of Assessment time, Group type** |  |  |  |  |  |
| Intercept | 0.167 | 0.240 | -0.303 – 0.637 | 0.698 | 0.485 |
| Assessment time | -0.016 | 0.037 | -0.036 – 0.057 | -0.426 | 0.670 |
| pMCI diagnosis | 4.497 | 0.379 | 3.754 – 5.241 | 11.855 | **<0.001** |
| Assessment time * pMCI diagnosis | 0.350 | 0.056 | 0.240 – 0.460 | 6.226 | **<0.001** |
| **Interaction of Assessment time, Group type, Gender** |  |  |  |  |  |
| Intercept | 0.208 | 0.370 | -0.517 – 0.932 | 0.562 | 0.574 |
| Assessment time | 0.010 | 0.055 | -0.098 – 0.118 | 0.181 | 0.857 |
| pMCI diagnosis | 4.680 | 0.602 | 3.500 – 5.860 | 7.773 | **<0.001** |
| Males | -0.069 | 0.489 | -1.027 – 0.888 | -0.142 | 0.887 |
| Assessment time * pMCI diagnosis | 0.554 | 0.083 | 0.391 – 0.717 | 6.659 | **<0.001** |
| Assessment time * Males | -0.046 | 0.072 | -0.187 – 0.095 | -0.640 | 0.522 |
| pMCI diagnosis * Males | -0.271 | 0.780 | -1.800 – 1.258 | -0.347 | 0.728 |
| Assessment time * pMCI diagnosis * Males | -0.346 | 0.108 | -0.558 – -0.134 | -3.202 | **0.001** |
| **Interaction of Assessment time, Group type, APOEε4** |  |  |  |  |  |
| Intercept | 0.072 | 0.261 | -0.440 – 0.585 | 0.276 | 0.782 |
| Assessment time | -0.027 | 0.042 | -0.109 – 0.055 | -0.640 | 0.522 |
| pMCI diagnosis | 4.323 | 0.416 | 3.508 – 5.137 | 10.399 | **<0.001** |
| APOEε4 carrier | 0.377 | 0.414 | -0.435 – 1.188 | 0.910 | 0.363 |
| Assessment time * pMCI diagnosis | 0.164 | 0.078 | 0.011 – 0.318 | 2.101 | **0.036** |
| Assessment time * APOEε4 carrier | 0.046 | 0.086 | -0.123 – 0.214 | 0.528 | 0.598 |
| Assessment time * pMCI diagnosis * APOEε4 carrier | 0.276 | 0.121 | 0.039 – 0.513 | 2.286 | **0.022** |

**Table 3.**
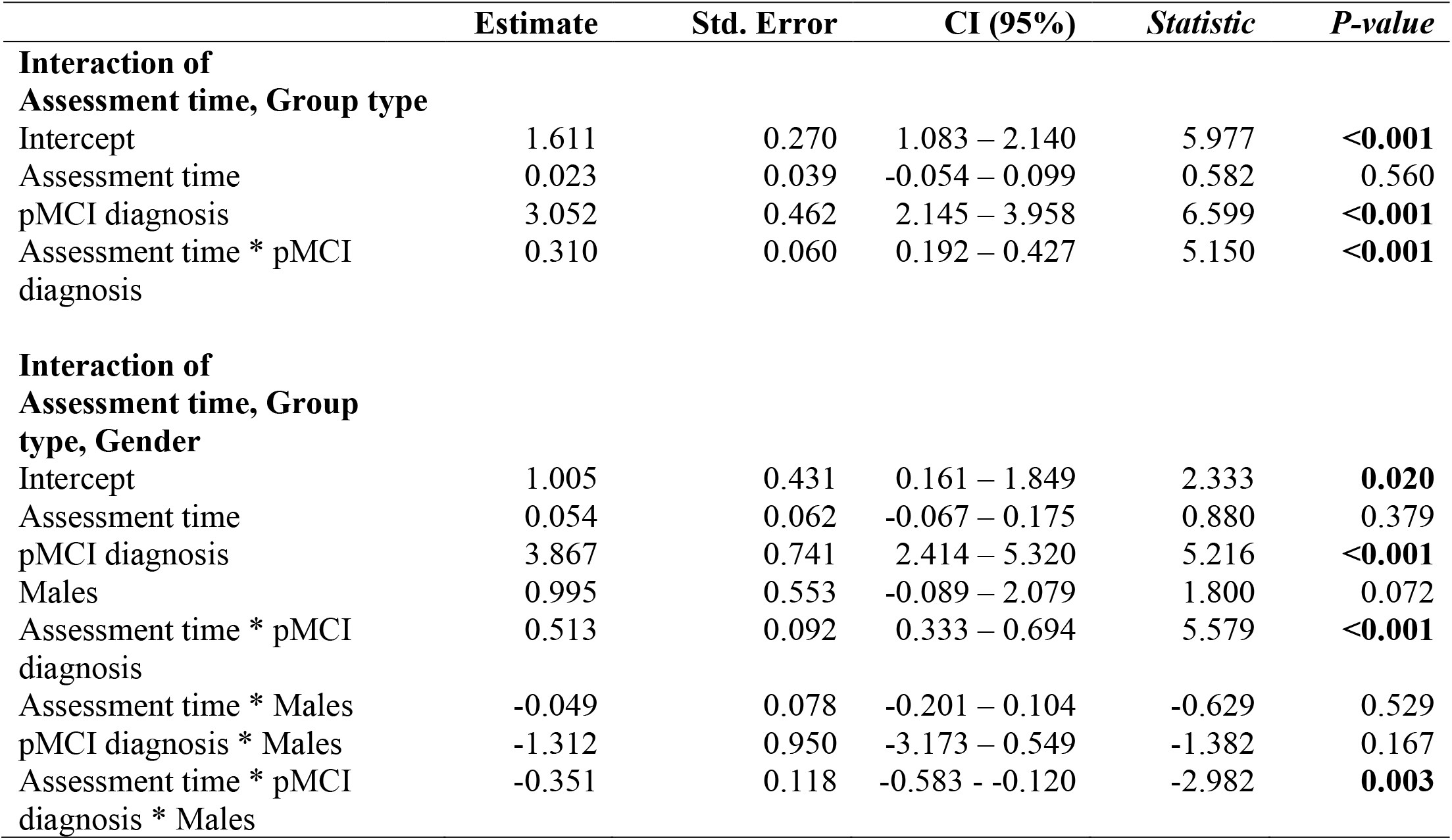
Multilevel modeling results for group type stable mild cognitive impairment (sMCI) compared to progressive mild cognitive impairment (pMCI). Bold p-values are <.05

Given the significant interaction of group type and assessment on the BAG, additional subject-level covariates were included in the multilevel models to test whether the difference in rate of BAG change over assessment across groups (i.e., the 2-way interaction effect) depended on subject-specific characteristics. These models examined the influence of gender (male or female) using the 3-way interaction of assessment, group type, and gender. Comparing pMCI to sCN found the difference in BAG rate of change was stronger for females compared to males (*b* = .346, *p* < .001), and similarly for pMCI compared to sMCI (*b* = .351, *p* = .003). Visualizing the trajectory of each group over assessments shows a slight increasing BAG trajectory for pMCI females compared to males across groups (Figure 4).

**Figure 4.**
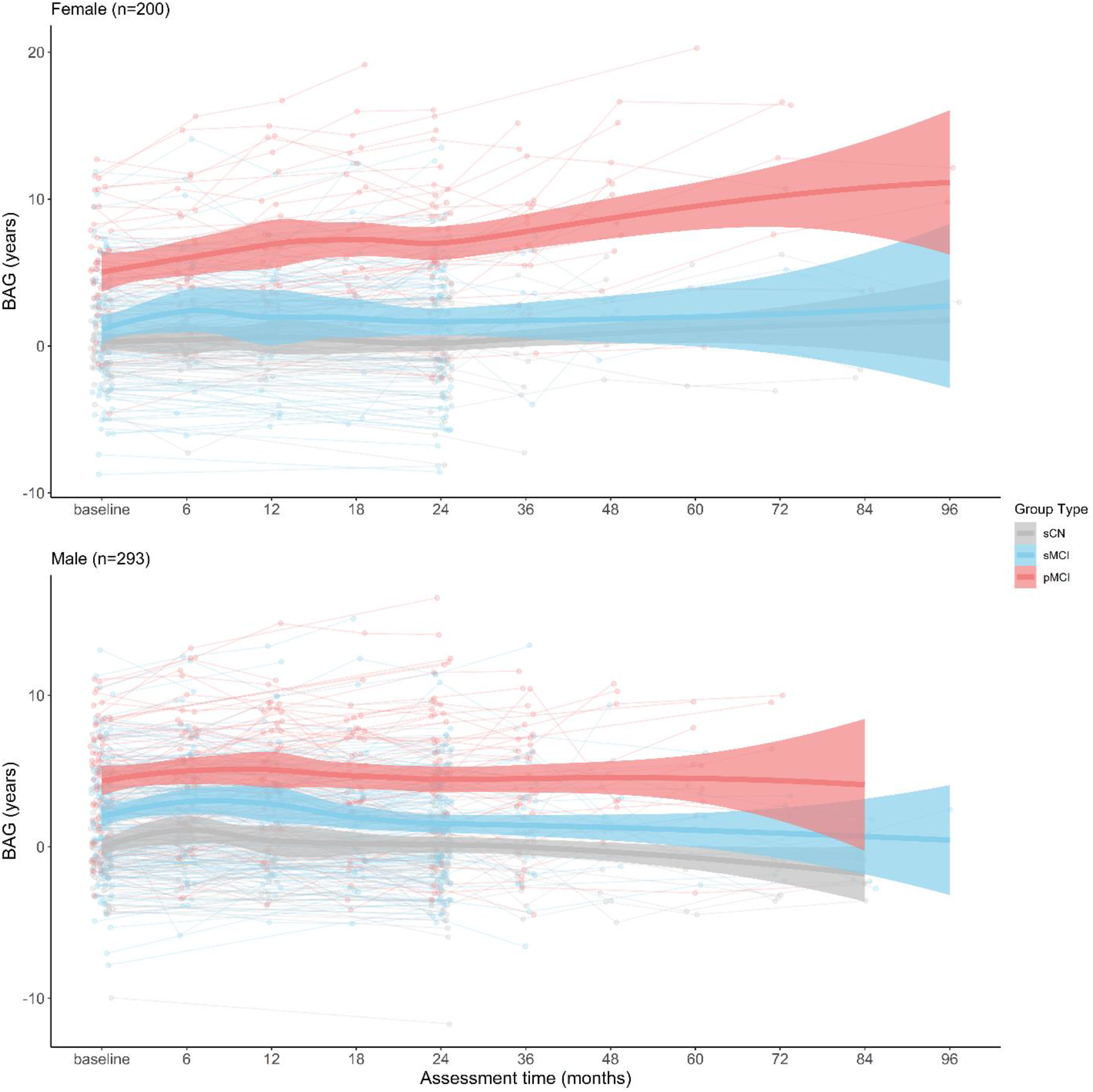
Individual and group brain age gap (BAG) values at each assessment month following baseline for females (top) and males (bottom). Thick lines represent BAG of all subjects for each group across assessments, fitted by loess curve. Shaded area along each curve indicates the 95% confidence interval. Individual dots and lines represent individual subjects for each group. Groups are given as stable cognitively normal (sCN), stable mild cognitive impairment (sMCI), and progressive mild cognitive impairment (pMCI). Assessment time represents the months following the baseline (initial) assessment.

APOEε4 carriership was examined similarly (i.e.., comparing carriers to non-carriers). The respective 3-way interaction was significant when comparing pMCI to sCN (*b* = .276, *p* = .022), but not when comparing pMCI to sMCI (*b* = .184, *p* = .128).

### Trajectory analysis of pMCI

The increasing change in BAG for the pMCI group was further examined using piecewise linear segments to assess whether the rate of increase in BAG was greater following the diagnosis of AD. Piecewise linear segments of preAD and postAD as fixed effects were best modeled when varying individuals over the preAD and postAD assessments in the random effects, compared to using only preAD (*X^2^*(3) = 28.534,*p* < .001) or only postAD(*X^2^*(3) = 9.362, *p* = 0.025). Using this random effect structure, the rate of increase in BAG was greater for the postAD segment (*b* = .280, *p* = .007) compared to the preAD segment (*b* = .209, *p* = .010).

To examine whether the rate of change in BAG differed significantly before and after AD diagnosis, multilevel modeling of the 2-way interaction of assessment and conversion variable (i.e. a binary variable indicating whether assessments were before and after AD) was conducted. This interaction was non-significant (*b* = .108, *p* = .286), suggesting the rate of increase in BAG did not differ significantly before and after conversion.

### Feature Importance

Top neuroimaging features at least one standard deviation above the average feature importance value were considered to contribute positively to the prediction of the BAG (Figure S1). Top MRI features of cortical thickness were the left inferior parietal cortex, inferior temporal gyrus, lingual gyrus, pars triangularis, precuneus cortex, rostral middle frontal gyrus, superior temporal gyrus, and right entorhinal cortex, lateral orbital frontal cortex, middle temporal gyrus, postcentral gyrus, superior frontal gyrus, and supramarginal gyrus. Top cortical FDG-PET features were bilateral fusiform gyrus, precuneus cortex, rostral middle frontal gyrus, and pars opercularis, as well as left caudal anterior-cingulate cortex, lingual gyrus, postcentral gyrus, superior temporal gyrus, transverse temporal cortex, and finally right inferior parietal cortex, lateral occipital cortex, pericalcarine cortex, superior parietal cortex, supramarginal gyrus, and insula. Top subcortical FDG-PET features included bilateral cerebellum cortex, left thalamus, and right pallidum and accumbens area.

## DISCUSSION

To our knowledge, the present study is the first to use a longitudinal data analysis approach to investigate the pattern and rate of change in the difference between brain age and chronological age (i.e. BAG) over time. We found that the estimated BAG values were not only consistently higher in MCI participants who progressed to AD compared to CN or MCI individuals who did not progress, but they also increased at a faster rate. Furthermore, this difference was moderated by gender, where the increase was larger in females. Additionally, the difference in BAG increase between MCI individuals who progressed and sCN was moderated by APOEε4 carriership where the increase was larger in APOEε4 carriers. Finally, investigation of the increasing rate of BAG for the pMCI group showed a greater slope across assessments following the diagnosis of AD compared to assessments preceding the AD diagnosis, however this increasing rate overall was not significantly moderated by timepoint of AD conversion. While the specific cause for the increasing rate of BAG change is unclear, these results demonstrate the utility of the BAG as a biomarker for understanding group specific temporal patterns related to AD progression.

Interestingly, the non-linear assessment time effect was not significant, though this is likely due to the test considering data from all groups. When comparing group effects, the BAG was observed to increase at a significantly faster rate for MCI individual who progressed compared to the stable individuals. Several cross-sectional studies have observed larger BAG in MCI individuals who progressed to AD [18,20]. However, these did not account for individual variances, which were considered in our multilevel models. Accounting for individual differences is important given expected differences in genetic and environmental effects across aging [23].

Within pMCI, we also found that BAG increased at a numerically greater rate after conversion to AD compared to before conversion, however this increase was not statistically significant. This is likely due to the small sample size for pMCI with more than one assessment with AD diagnosis (*N*=47). Our findings suggest that, although the progression of AD-related brain changes likely precede clinical diagnosis, there may be a change point for when the BAG begins to accelerate. Identifying this change point could potentially create a more personalized prediction of AD progression. Future studies with larger longitudinal samples are warranted.

How covariates related to AD progression moderate the BAG trajectory are also important to consider. Women are afflicted with AD at a greater rate than men after accounting for greater life expectancy [51], and may show a greater rate of clinical decline if afflicted [52]. The reasons are disputed but may include neurological consequences of menopause triggering pathological protein aggregation in women [53–55], or differential effects of APOEε4 [56]. Our study contributes to this growing literature, reporting that group differences in the rates of BAG increase were greater for female pMCI than for males.

Similarly, we found rates of BAG increase were greater for pMCI APOEε4 carriers than for non-carriers, which is in line with a recent study (unrelated to the BAG) which found carriers of APOEε4 were associated with faster progression of AD-related dementia [57]. Taken together with results from the model including sex, these results are particularly interesting as volume in the hippocampus, a region implicated in AD pathology [58], has previously been found to be significantly reduced in female APOEε4 carriers with MCI, compared to female non-APOEε4 carriers with MCI [59]. Further, APOEε4 has been shown to improve accuracy for predicting AD conversion [21] and interact with race and sex. White and black adult APOEε4 carriers were shown to have overlapping but differential cognitive resilience factors [60], a predictor for AD risk (though our sample size limited an investigation into race). Amyloid-β deposition may also have moderating effects on the BAG trajectory, given frequent associations between amyloid and AD [2]. Indeed, a recent study has shown the brain age prediction framework can be used to distinguish between amyloid-β negative or positive status [61]. The present study did not consider this covariate due to sample size constraints, but importantly, the mentioned study could only significantly distinguish between the two statuses when increasing the sample size beyond only ADNI, which introduces potential variability in site effects [62]. Clearly, the moderating effects of covariates such as sex, race, Amyloid-β, and APOEε4 (among others) on the BAG trajectory is complex but necessary to evaluate. Future studies using a longitudinal design to study the BAG’s temporal pattern should consider these covariates.

Taken together, our results are line with the larger literature of AD-related changes in the brain suggesting accelerated aging, such as gray matter and cortical thickness atrophy [6,19,63]. While normal aging is associated with gray matter atrophy between 0.2-0.5% annually, longitudinal MRI studies have shown annual atrophy rates of 2-3% in AD patients [64]. However, reports of differential cortical thickness changes across regions of the brain [65] suggests the utility of the BAG as a biomarker of brain changes could be enhanced through improved brain parcellation.

While the present paper quantified importance values of brain regions for the SVR’s model performance, these values were based on sCN neuroimaging data, and thus represent the association between improved prediction performance and aging in the sCN group and not necessarily the effect of AD-related atrophy. However, regions we determined as ‘important’ have previously been identified, where uptake values from FDG-PET in regions including the anterior cingulate cortex and precuneus [66], and left superior temporal gyrus, and right insula [67] were negatively correlated with age in CN patients. Similar regions have also been observed in AD patients to show gray matter atrophy, such as the precuneus and inferior temporal, superior frontal, supramarginal, and lingual gyri, as well as decreased cerebral blood flow in the anterior cingulate, right insula, and precuneus, as well as superior temporal, orbital frontal, lingual, and fusiform gyri [68–70]. If BAG demonstrates value as a biomarker for AD, regions which improve its prediction are likely associated with AD. However, the direct relationship of these regions to AD and their causal effect remains to be investigated.

Nevertheless, the results of the present study have potential clinical relevance. Previous studies have already demonstrated not only an association between a positive BAG and increased likelihood of AD, but also a number of other neurological conditions [11]. Our findings expand on these studies by considering a longitudinal design and how related covariates may influence the trajectory of the BAG. Longitudinal designs are critical for early detection of neurodegenerative diseases.

Limitations to this study should also be considered. First, the SVR model was trained using only the ADNI dataset and does not necessarily generalize to other datasets with different racial or socioeconomic distributions, or methodological differences. The advantage of a single dataset is that the results can be directly compared to other studies working with the ADNI, and differences would not be attributable to variations in dataset or methodology. Related to all PET imaging data (including from ADNI) are partial volume effects due to low resolution, which may cause the activity from small ROIs to be underestimated [71]. On the other hand, whether correction methods for these effects are reliable or have a noticeable impact on results are controversial [72–74]. Future brain aging studies using FDG-PET may wish to consider these effects.

Additionally, other machine learning methods must be considered. We compared relevance vector regression (RVR) [7] and LASSO [75] with the SVR presented here achieving the best performance (comparisons not shown). Although optimization in the SVR lead to more accurate representation of the BAG trajectories, its computational cost was high. RVR has the potential for clinical adoption because of its low computational cost from not requiring parameter optimization. Other methods such as deep neural networks have the potential to improve BAG estimation as well [9,76].

## CONCLUSION

Our study contributes to the existing literature by taking a multimodal, longitudinal approach to examine the temporal patterns of brain aging and found that brain aging occurs at an accelerated rate for those with pMCI compared to stable individuals. It further suggests that there may be a point of acceleration, although this finding needs replication in a larger longitudinal sample. These dynamic changes as subjects progress from MCI to AD are further moderated by both gender and APOε4 status. Describing the temporal trajectory for brain aging is particularly valuable for understanding AD progression and improving early detection through predictive models. Additionally, BAG prediction performance was improved using both MRI and FDG-PET data, suggesting complementary neuroimaging measures should be considered in BAG studies. Future studies may further examine the influence of other covariates on the BAG, explore individualized change points in pMCI trajectory, and expand the generalizability of the BAG machine learning models.

