## Supplementary Materials for "Investigating the temporal pattern of neuroimaging-based brain age estimation as a biomarker for Alzheimer’s Disease related neurodegeneration"

TABLE S1

| 'lh_bankssts' | 'lh_superiorparietal' | 'rh_precentral' |
| --- | --- | --- |
| 'lh_caudalanteriorcingulate' | 'lh_superiortemporal' | 'rh_precuneus' |
| 'lh_caudalmiddlefrontal' | 'lh_supramarginal' | 'rh_rostralanteriorcingulate' |
| 'lh_cuneus' | 'lh_temporalpole' | 'rh_rostralmiddlefrontal' |
| 'lh_entorhinal' | 'lh_transversetemporal' | 'rh_superiorfrontal' |
| 'lh_frontalpole' | 'rh_bankssts' | 'rh_superiorparietal' |
| 'lh_fusiform' | 'rh_caudalanteriorcingulate' | 'rh_superiortemporal' |
| 'lh_inferiorparietal' | 'rh_caudalmiddlefrontal' | 'rh_supramarginal' |
| 'lh_inferiortemporal' | 'rh_cuneus' | 'rh_temporalpole' |
| 'lh_insula' | 'rh_entorhinal' | 'rh_transversetemporal' |
| 'lh_isthmuscingulate' | 'rh_frontalpole' | Left_Cerebellum_Cortex |
| 'lh_lateraloccipital' | 'rh_fusiform' | Left_Thalamus_Proper |
| 'lh_lateralorbitofrontal' | 'rh_inferiorparietal' | Left_Caudate |
| 'lh_lingual' | 'rh_inferiortemporal' | Left_Putamen |
| 'lh_medialorbitofrontal' | 'rh_insula' | Left_Pallidum |
| 'lh_middletemporal' | 'rh_isthmuscingulate' | Brain_Stem |
| 'lh_paracentral' | 'rh_lateraloccipital' | Left_Hippocampus |
| 'lh_parahippocampal' | 'rh_lateralorbitofrontal' | Left_Amygdala |
| 'lh_parsopercularis' | 'rh_lingual' | Left_Accumbens_area |
| 'lh_parsorbitalis' | 'rh_medialorbitofrontal' | Right_Cerebellum_Cortex |
| 'lh_parstriangularis' | 'rh_middletemporal' | Right_Thalamus_Proper |
| 'lh_pericalcarine' | 'rh_paracentral' | Right_Caudate |
| 'lh_postcentral' | 'rh_parahippocampal' | Right_Putamen |
| 'lh_posteriorcingulate' | 'rh_parsopercularis' | Right_Pallidum |
| 'lh_precentral' | 'rh_parsorbitalis' | Right_Hippocampus |
| 'lh_precuneus' | 'rh_parstriangularis' | Right_Amygdala |
| 'lh_rostralanteriorcingulate' | 'rh_pericalcarine' | Right_Accumbens_area |
| 'lh_rostralmiddlefrontal' | 'rh_postcentral' |  |
| 'lh_superiorfrontal' | 'rh_posteriorcingulate' | |

85 cortical and subcortical regions of interest. MRI and FDG-PET features were parcellated based on these regions. “lh” represents left hemisphere, and “rh” represents right hemisphere.

TABLE S2

| **Patch Size** | **SVR Kernel** | **RMSE** | **Lower CI** | **Upper CI** |
| --- | --- | --- | --- | --- |
| 500 | Linear | 4.85 | 4.83 | 4.86 |
|  | Polynomial | 5.15 | 5.13 | 5.16 |
| 1000 | Linear | 4.98 | 4.97 | 5.00 |
|  | Polynomial | 4.97 | 4.96 | 4.99 |
| 2000 | Linear | 4.79 | 4.77 | 4.81 |
|  | Polynomial | 5.10 | 5.08 | 5.12 |
| 5000 | Linear | 5.17 | 5.16 | 5.19 |
|  | Polynomial | 5.05 | 5.03 | 5.07 |
| 10000 | Linear | 5.16 | 5.14 | 5.17 |
|  | Polynomial | 5.10 | 5.08 | 5.11 |

Comparison of support vector regression (SVR) root mean squared error (RMSE) results for different neuroimaging patch sizes (voxels/patch). Confidence intervals (CI) are based on 1000 bootstrap repetitions of training the model.

FIGURE S1

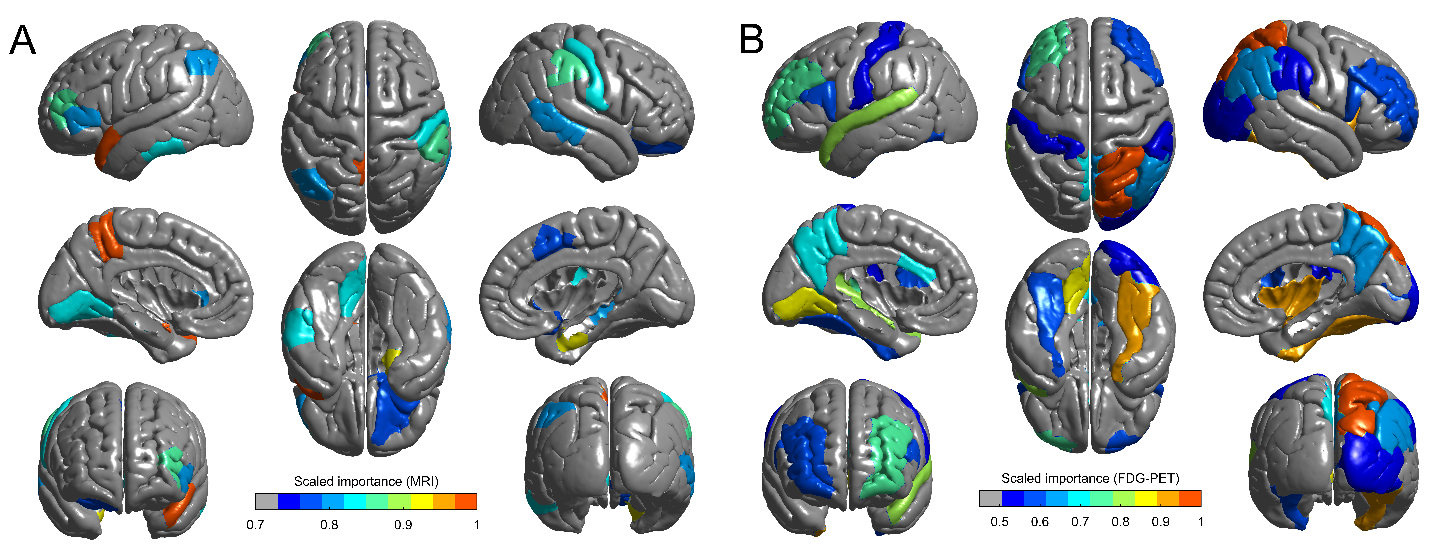

Min/max scaled importance values for top (greater than 1 SD from the mean) (A) MRI and (B) cortical FDG-PET features. Importance values represent the ratio of MAE after removing that feature, to the original MAE with all features, where a higher value indicates a greater feature importance for prediction.
